# Optical and magnetic properties of particles containing guanine in strong basic solution

**DOI:** 10.1101/2021.03.22.436526

**Authors:** Masakazu Iwasaka

**Affiliations:** Hiroshima University Kagamiyama 1-4-2, Higashihiroshima, Hiroshima 739-8527 Japan

## Abstract

Guanine based particle is one of very efficient material to control light especially near the surface of animal body. Past studies reported that platelets of anhydrous guanin crystal are utilized by living creatures, and they can change their colour and light reflection intensity when the arrangement of platelets. Guanine has relatively high reflective index and its particle can exhibit a unique optical property. Under higher pH which can be provided by a high concentration of NaOH aqueous solution, guanine molecules formed a sodium salt particle which contain guanine. Here this work presents that particles containing guanine in basic aqueous solution with NaOH exhibited very strong shining and light reflection switching by magnetic fields. In addition, adding an extracted solution of a fish iridophore enhanced the formation of intensively light reflecting guanine particles floating in a strong basic solution.

## Introduction

In a living tissue, a gradient in density of tissue materials may have a role to control a biophysical function. Especially a periodical displacement of materials with different vale of refractive index can cause an interference of electromagnetic waves in them, and in cases of visible light, the light being reflected in multiple layers with different refractive indexes perform a modulation of colour and light intensity.^1–4^ This mechanism was discussed on huge numbers of studies dealing with animal body surface display as structural colours.^5–10^ In addition, recent studies are struggling with a rapid development and opening new horizon in interdisciplinary fields of biology, biological material sciences, optics & photonics and engineering.^11–15^

One of the main topics of biologically designed light control material and structure is an organic micro particle made of guanine.^16–21^The biogenically crystallized platelets are frequently observed in a thin tissue of fish body skin^16,17,19^, eye^19,21^ and light emitting photophores^18,20^. Previous studies concerning biogenic guanine crystals from animals such as fish suggested that biogenic guanine crystals have a special optical property to enhance shining on its surface. Guanine has an anisotropy in refractive index, and a multiple layer of guanine platelets exhibits a broadband reflection which is the mechanism for the silver shining on the skin of fishes.^22^ The crystal platelets of guanine exhibited a clear response to external magnetic fields.^23–26^It was found that goldfish guanine platelet can be controlled its orientation and a specific angle of the micro-mirror reflecting light intensely was determined.^23,24^ The magnetic orientation mechanism was theoretically explained by diamagnetic anisotropy of the platelet. Furthermore, applying high magnetic field at 10 Tesla induced a structural colour change in a guanine platelets aggregation on a goldfish scale.^26^

A periodical structure in living tissue repeating with a distance equal to light wavelength can act as photonic crystal, as well as the photonic crystals made of solid state materials such as silicon. Several kinds of crystallographic studies were reported in order to clarify the reason for the silver shining in the biogenic guanine crystals.^27–29^ In addition, artificially synthesized guanine crystals were investigated by several groups.^28–30^ But little knowledge was obtained from the view point of light reflectivity of the synthesized guanine crystals or guanine particles. The present study is designed to explore an intensively light reflecting condition in a higher basic solution system with dissolved or colloidal suspension of guanine.

Among the previous studies developing synthetic guanine crystals, Oaki et al.^29^ succeeded in forming platelet-like crystals by using ammonia solution and additives, and Gur et al.^28^ discovered a novel method to induce anhydrous, mono-hydrate guanine crystals or guanine sodium salt by changing pH. Although these studies provided detailed knowledges for the crystallographic properties, checking the light control function of the synthesized crystals were remained. Also, in a re-crystallization work of the author’s group,^30^ recrystalized guanine particles in an aqueous solution at neutral pH after cooling down from 100°C did not so intense reflection.

At present it seems that no work is reaching the stage of realizing the same intense reflection as biogenic guanine platelets in artificially synthesized guanine crystals or particles. However, the present report shows that in a transient process of precipitation of guanine in a strong basic solution, effects of inhomogeneity of solvent have an important role in enhancing the reflectivity of particles containing guanine. In addition, for the first time it is found that an extraction from iridophore (Iridophore extraction, IE) increase the reflectivity.

## Results and Discussion

A condensed solution of guanine in a strong basic solution of NaOH was prepared in the first, as shown in Method section. The solution was a kind of colloidal suspension of fine guanine particles floating in the basic solution. For the initiation of precipitation of guanine particles, this work employed a “trigger solution,” which was a mixture of NaOH solution and HCl. The trigger solution at pH of around 12.5 containing NaCl was added to the condensed guanine colloidal suspension at pH of over 13. As shown in Figure 1, guanine fine particles (white, clouded part) once disappeared and formed light reflecting particles (40min), then after the 2^nd^ stimulation with the iridophore extraction, more brilliant particles were generated at 92min.

**Figure 1.**
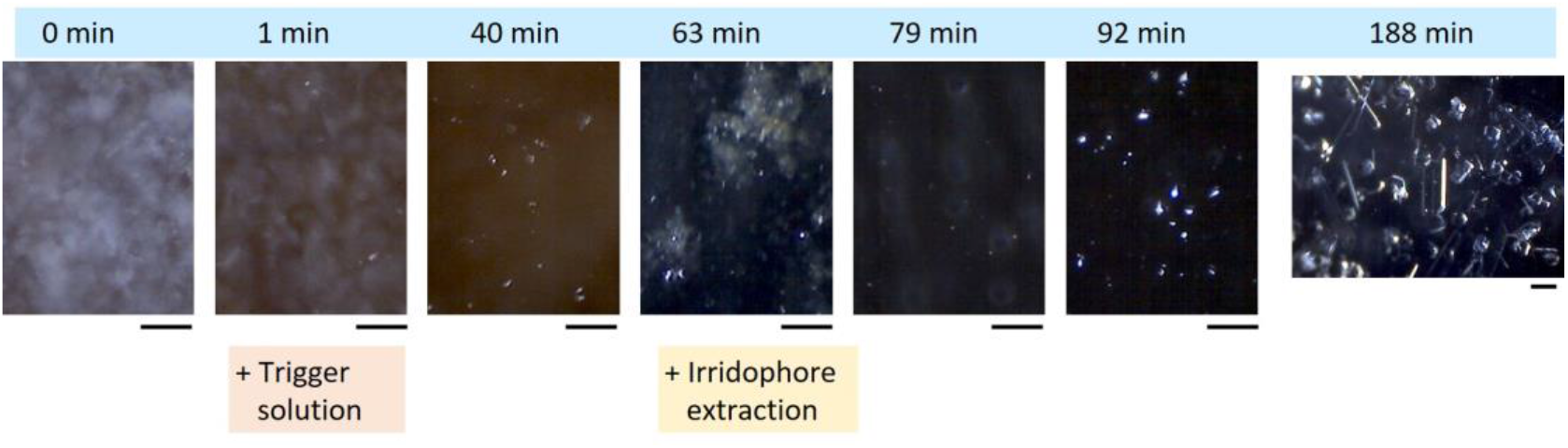
Time course of change in light reflection behaviours of condensed guanine suspension at high pH (>13). Two stimulations with a trigger solution and an iridophore extraction (from carp fish) was added at 1min and 63min, respectively. Bar is 200μm.

The light illumination was provided from side (dark-field illumination) at an incident angle of 75°to air-liquid boundary. In this case, the suspension was set in a well and the re-crystallization (precipitation) proceeded without continuous stirring (Figure 8a).

It was suggested that adding the extracted constituents from carp fish skin, mainly from iridophore, induced the enhancement of light spot intensity of the precipitated particles. At 63min, there were some white particles. These particles were fish guanine crystal platelets contained in the added IE, but in 79min these white particles disappeared and not remained in the suspension as original platelets. The brilliant white particles were newly generated in the suspension with IE.

Exposing the sample to the air continuously and vaporizing water transformed these particles showing intense light spots to platelets, which were speculated to be a guanine sodium salt in strong basic solution. The platelets settled on the bottom of well, and were not floating and reflected light only at edges of platelet.

Next, guanine precipitation was accelerated by stirring the suspension on a glass substrate (Figure 2), and I compared the effects of trigger solution only and both the trigger solution and IE of goldfish. An example image of IE is shown in Figure 2b. The solution contained materials attaching to the scales which were detached from goldfish body. As well as IE extract, guanine platelets, chromatophore pigments and cellular components were mixed. In Figure 2a, Left-top image is the right after mixing a trigger solution into a strong basic solution with high density of guanine. Only with the trigger solution, intermittent stirrings brought platelet-like crystals which sedimented on bottom (right-top image) although part of the platelets showed a brilliance similar to that of biogenic platelet (Figure 2c). On the other hand, as shown in bottom-middle image, after 15min from adding both trigger solution and iridophore extraction, very brilliant, floating and flickering particles appeared.

**Figure 2.**
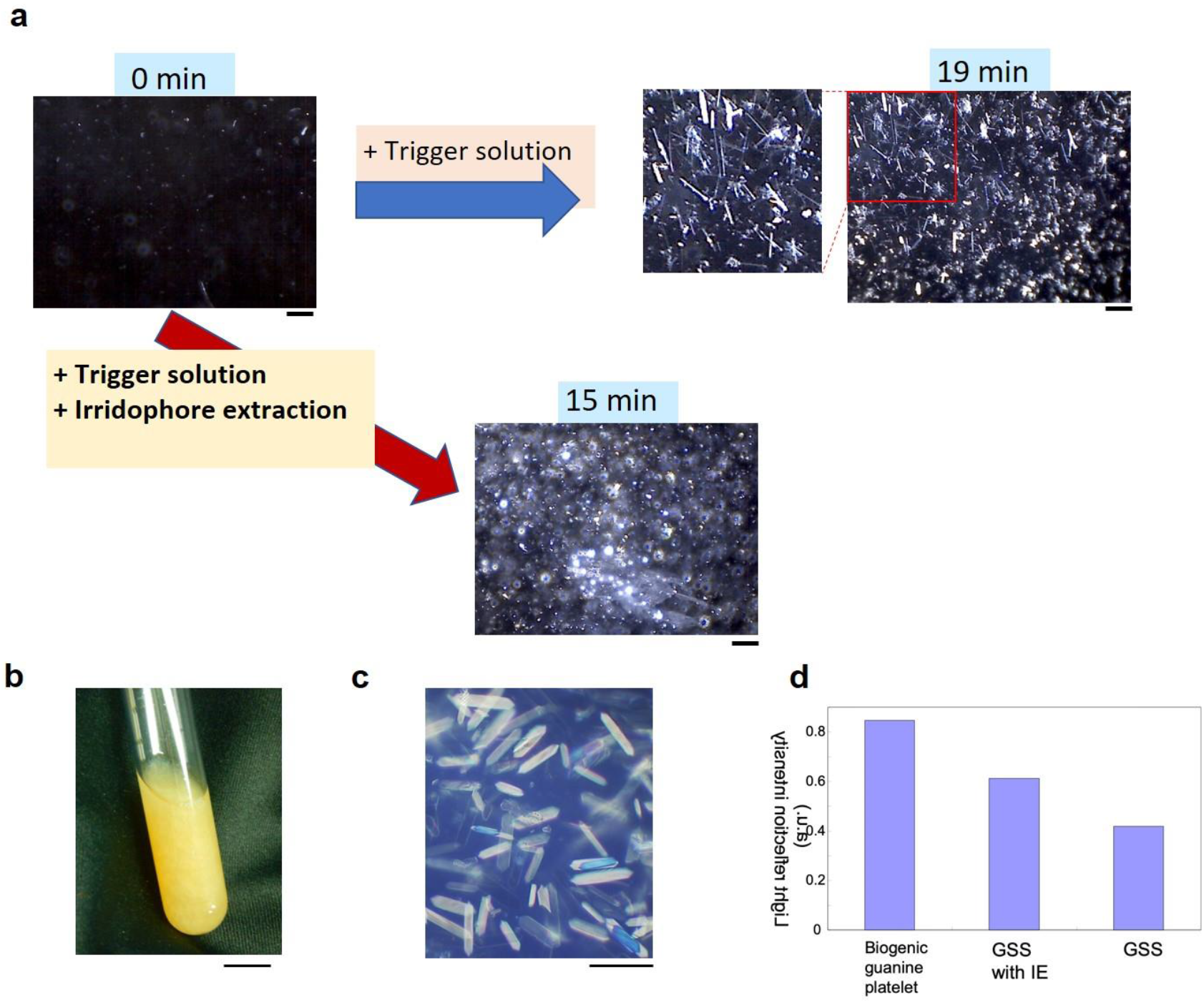
Comparison of guanine sodium salt particles formed under different stimuli with biogenic guanine platelets from goldfish. **a**. Particle formation in a thin layer of mixture on glass substrate with stirring. Left-top image is the right after mixing a trigger solution into a strong basic solution with high density of guanine. Right-top image; only with trigger solution. Bottom-middle image; 15min after adding both trigger solution and iridophore extraction. Bar is 200μm. **b**. An example of goldfish iridophore extraction containing guanine platelets. Bar is 10 mm. **c**. Guanine crystal platelets of goldfish. Bar is 40μm. **d**. Light reflection intensity of three kinds of particles containing guanine, biogenic, guanine sodium salt (GSS) with iIE and GSS.

Figure 2d shows a quantitative analysis of the light spot intensity of three kinds of particles. The data was obtained by a brightness analysis under the same experimental setting for light illumination. Each of image was analysed to measure grey value (light spot intensity), and area occupied by particles was calculated, then the ratio of grey value and area was obtained. Biogenic guanine platelets exhibited strongest intensity, and the light spot of guanine sodium salt (GSS) with IE was more brilliant than GSS without IE.

To test the possibility that remaining biogenic guanine crystal platelets are still acting as brilliant particles in the strong basic solution at pH of 12 to around 13, following experiments were carried out. To an iridophore extraction suspension (containing 5 × 10^7^ /ml of guanine platelets), a strong basic solution at pH~12.5 was added, and a change in the shape of floating objects were observed under microscope in real time. Our main interest is the reflectivity and brilliance of the guanine particles under side light illumination. The biogenic particles having intense light spot (image of Figure 3a) under the illumination remained only air bubbles reflecting light at higher incident angle on its surface. The intensely reflecting biogenic particles were disintegrated. A bright field illumination image (Figure 3b) indicated the presence of constituents from chromatophore including iridophore in the strong basic solution.

**Figure 3.**
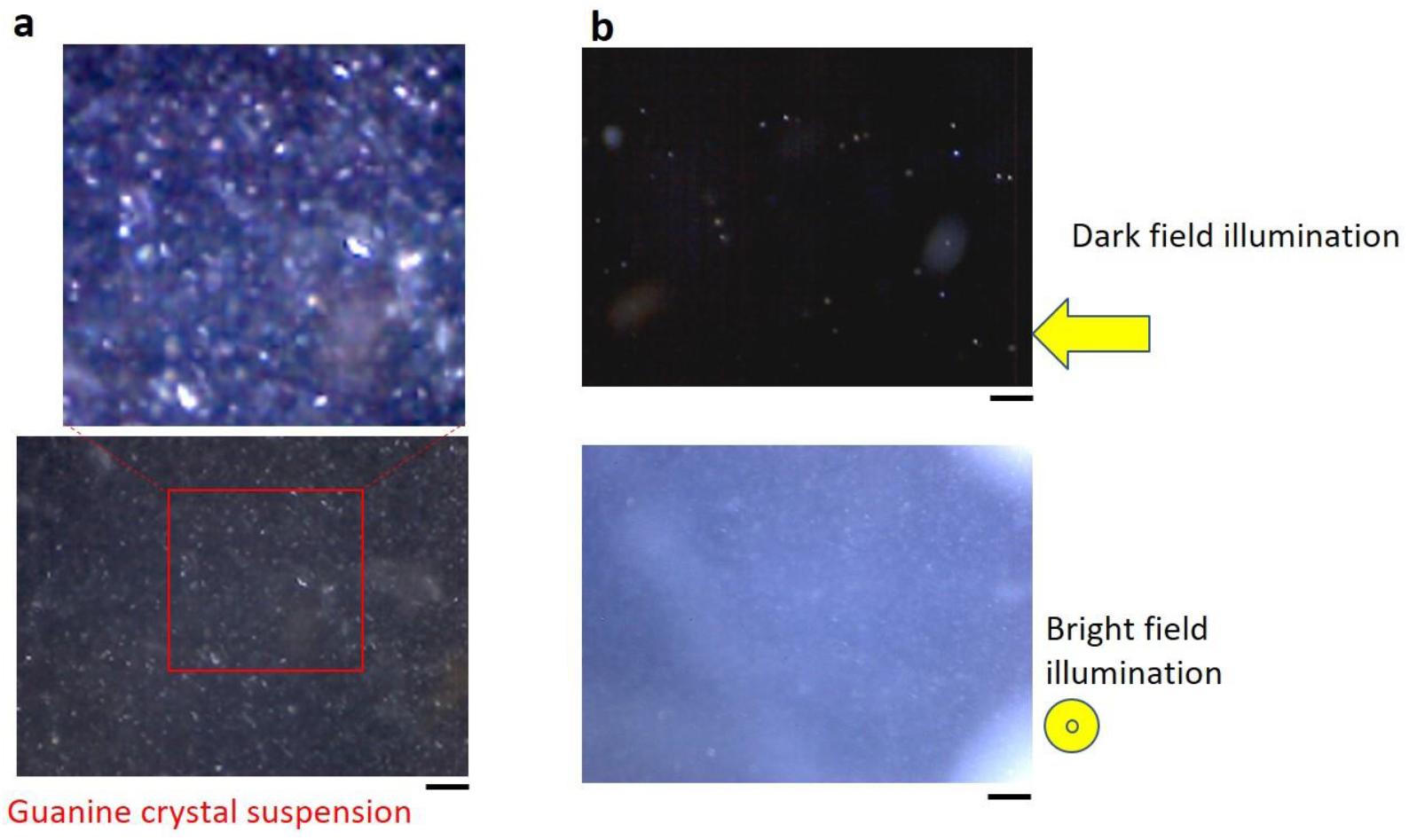
Dissolution of biogenic guanine platelets in iridophore extraction in the mixture with strong basic solution at pH~12.5. Dark field and bright field image of mixture containing dissolved guanine particles and iridophore extraction (lipid, protein, et al.). **a**. Before mixing. **b**. After mixing. Bar is 200μm.

Next this mixture of IE and strong basic solution was applied into triggered solution (Figure 4). Image taken in 36min have many brilliant and shining spots in a clouded background.

**Figure 4.**
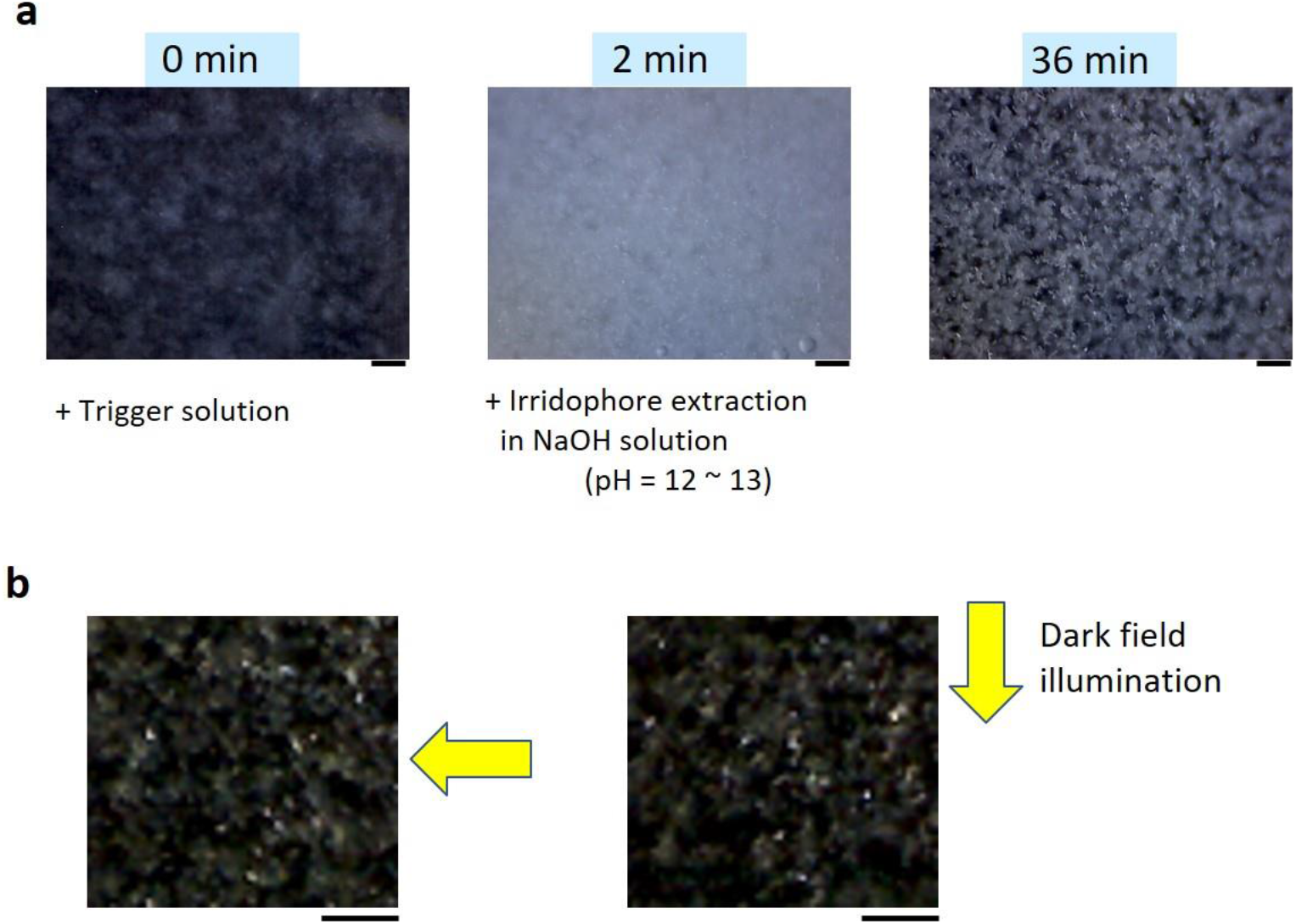
Effect of dissolved biogenic guanine platelets adding to triggered re-crystallization process. **a**. Time course of the mixing. **b**. Intensely light reflecting particles appeared in sodium salt precipitation in 40min, showing a light reflection anisotropy.

The position of the spots shifted when the direction of incident light by a LED hand light was changed. Our previous study revealed the light reflection anisotropy of fish guanine crystal platelets. The observed light spot shift means that we can generate a shining guanine particle by the action of the mixture of IE and strong basic solution.

A dependence of iridophore extraction density on light reflection intensity of guanine sodium salt particles was analysed in two samples. One of the IE solution was diluted 5 times by water. The diluted suspension (Figure 5b) generated smaller number of shining particles compared to the case with non-diluted IE (Figure 5a). An image analysis was carried out on these two images by comparing irradiance (grey value) at shining spot and surrounding dark background (Figure 5c). The averaged brightness of shining spots was more intense in the particles formed with dense (non-diluted) IE. And the difference of brightness between a particle (GP) and surrounding background (BG) was also distinct in the image-a with dense IE. This result indicates the role of a material which was involved in an IE or fish skin was essential for enhancing light spot intensity of particles containing guanine.

**Figure 5.**
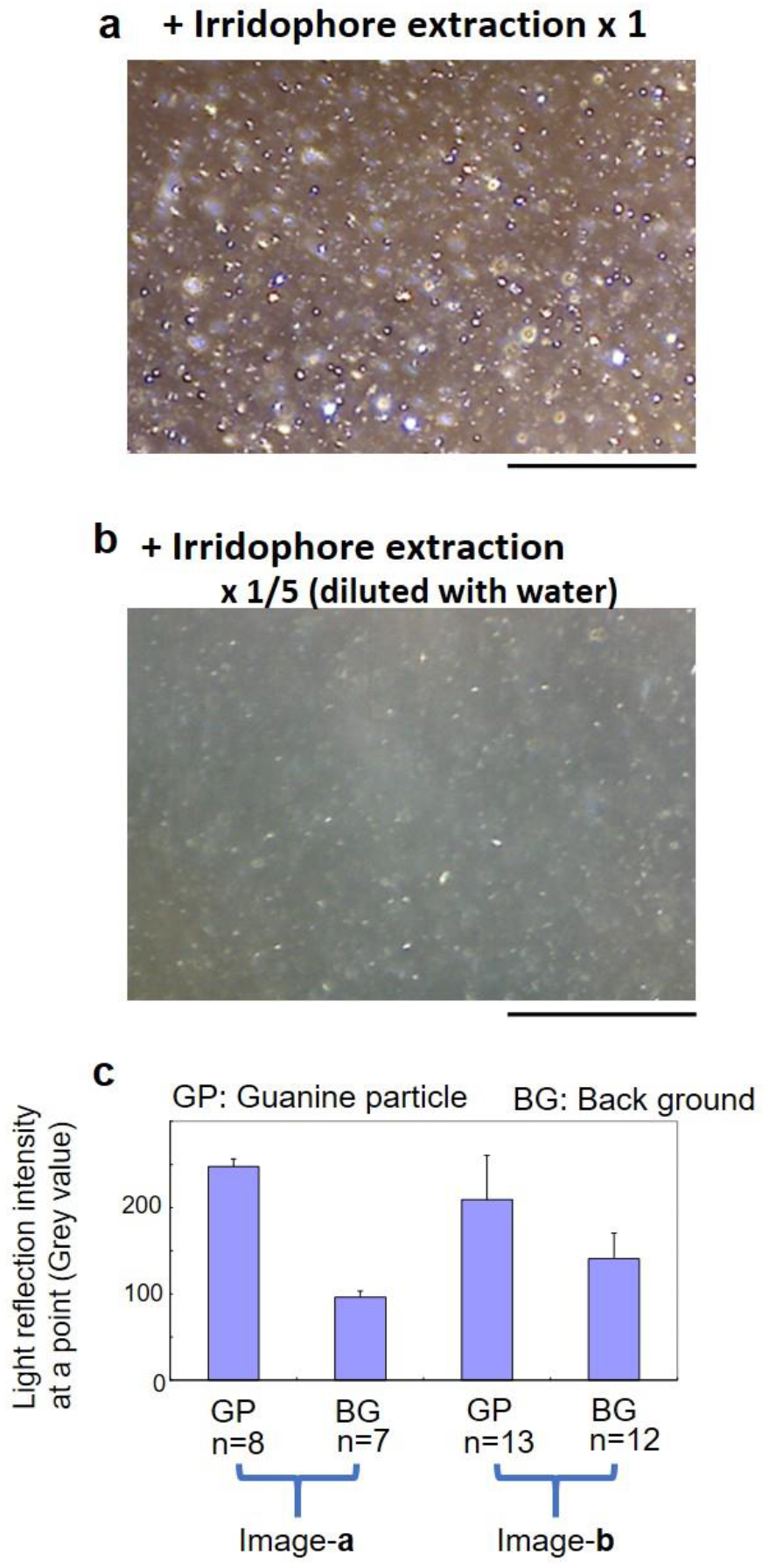
Effect of iridophore extraction (IE) density of the light reflecting guanine sodium salt particles. **a**. With addition of non-diluted IE. **b**. With addition of five times diluted IE. **c**. Analysis of the brightness in a particle (GP) and surrounding background (BG). Two sets of columns were obtained in the analyses on images of **a** and **b**. Bar is 500μm.

In Figure 5a, the precipitated guanine particles in a large size (~50 mm) and showed the same level of reflection intensity with fish guanine, while in figure5b, relatively small particles less than 1mm in length were dispersed between larger particles. The light randomly scattered in the area of smaller particles, and this condition decreased the contrast of particles in background in Figure 5b.

Final stage of present report is testing magnetic response of the developed particles containing guanine. Figure 6 shows a time course of the scattered light intensity from a glass tube containing a strong basic suspension of the precipitated guanine particles, which were detected by an optical fibre directing its axis parallel to the applied magnetic field at 300 mT. The incident light came from side (in perpendicular to the fibre axis). The experimental data shows that by magnetic orientation of guanine sodium salt particles floating in the solution, the detected light intensity decreased by the exposure of magnetic field at 300 mT. As shown in the model (bottom illustration) of Figure 6, it can be explained that a platelet-like particle can exhibit the light scattering intensity change under this configuration of incident light and detection. For this requirement, it was hypothesized that the platelet-like particle has guanine molecules aligning in parallel to the broadest surface of the model platelet from the consideration of diamagnetic anisotropy in guanine molecules assembly.

**Figure 6.**
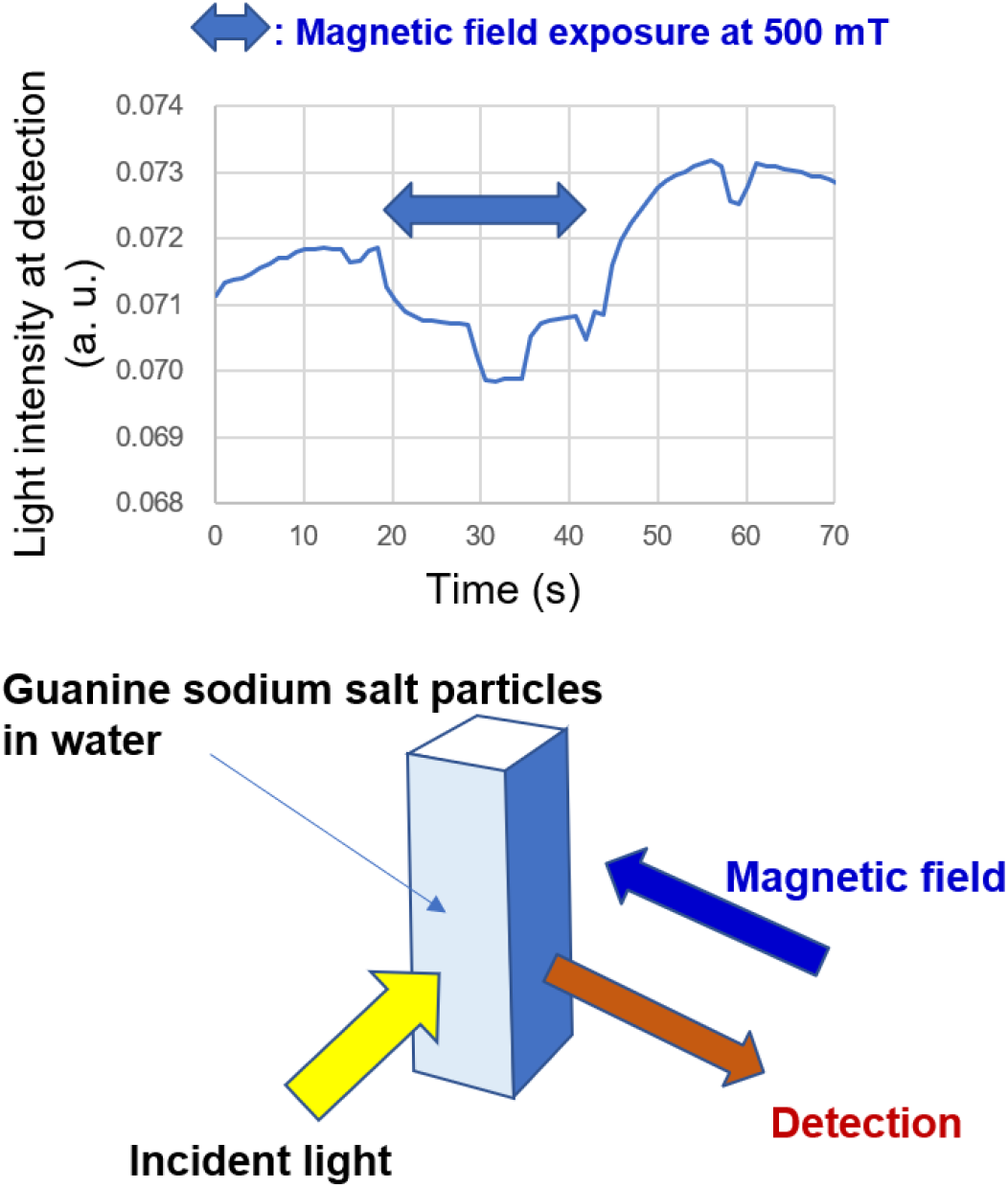
Magnetic orientation of guanine sodium salt particles and light reflection changes which were induced by the exposure of magnetic field at 300 mT.

Despite of this speculation concerning crystallographic aspect, we can conjecture that the developed intensely shining particles in a strong basic solution are owing its light control mechanism to an aggregation of guanine molecules. The investigation of the structure of the guanine aggregation is remained for other or future studies. If a platelet under high pH have a layered structure of guanine molecules, possibly the platelet has the similar diamagnetic anisotropy with the biogenic guanine crystals from goldfish’s iridophore. As shown in Figure 5a, the precipitated nano-/micro- platelets floating in a strong alkali liquid exhibited a very strong reflection of light.

We can hypothesize that if a platelet particle appeared in solution it is composed of layered structures of guanine molecular sheets. A previous study by Gur et al. indicated that the re-crystalized guanine in high pH sodium hydroxide became sodium crystals. Also, the present method has a demerit to form much of sodium chloride crystals which is surrounding the formed guanine platelets. At present, changing the pH of solution to neutral avoiding a degradation of recrystallized particles is hard to achieve. A rapid pH jump from pH > 13 to pH~7 sometimes remained very small amount of the particles but it reproducibility is small. As an additional succeeded result, the precipitated guanine particles showed properties which resembled those of biogenic guanine platelets in the magnetic response and light scattering.

Perhaps the aggregation can be modified by a constituent in iridophore extraction or fish skin tissue. Determining the detailed materials of the constituent may contribute to develop more shining and stable particles of guanine that can be called a mimetics of biogenic fish guanine platelets.

## >Conclusion

Under higher pH (>13), guanine molecules formed a shining salt particle which was resembled to the biogenic fish guanine platelets. The particles in strong basic aqueous solution exhibited a light reflection switching by magnetic fields. By adding an extracted solution of a fish iridophore to the re-crystallization process, it enhanced the formation of intensively light reflecting guanine particles.

## Methods

### Protocols for precipitation

Figure 7 shows main stream for obtaining guanine particle precipitation from a saturated guanine solution (solution S), which was prepared by dissolving guanine to strong basic solution whose pH was adjusted more than 13.9 by adding NaOH to a distilled water. The solution was prepared in a centrifugal tube (50ml, Iwaki Glass corp. Tokyo, Japan) and undissolved guanine were sedimented in the bottom of the tube, then the saturated guanine solution was in a colloidal clouded suspension where fine particles of undissolved guanine were floating. Initiation of precipitation (or re-crystallization) was done by adding one or two stimulation solution. One was a Trigger solution and the other was iridophore extraction solution. Trigger solution (solution T) was a mixture of NaOH aqueous solution whose pH was more than 13.9 and HCl. pH of the mixture was adjusted to 12.5~13.6. Ratio of the solution volume, S: T was set to be 10: 1 to 5: 1.

**Figure 7.**
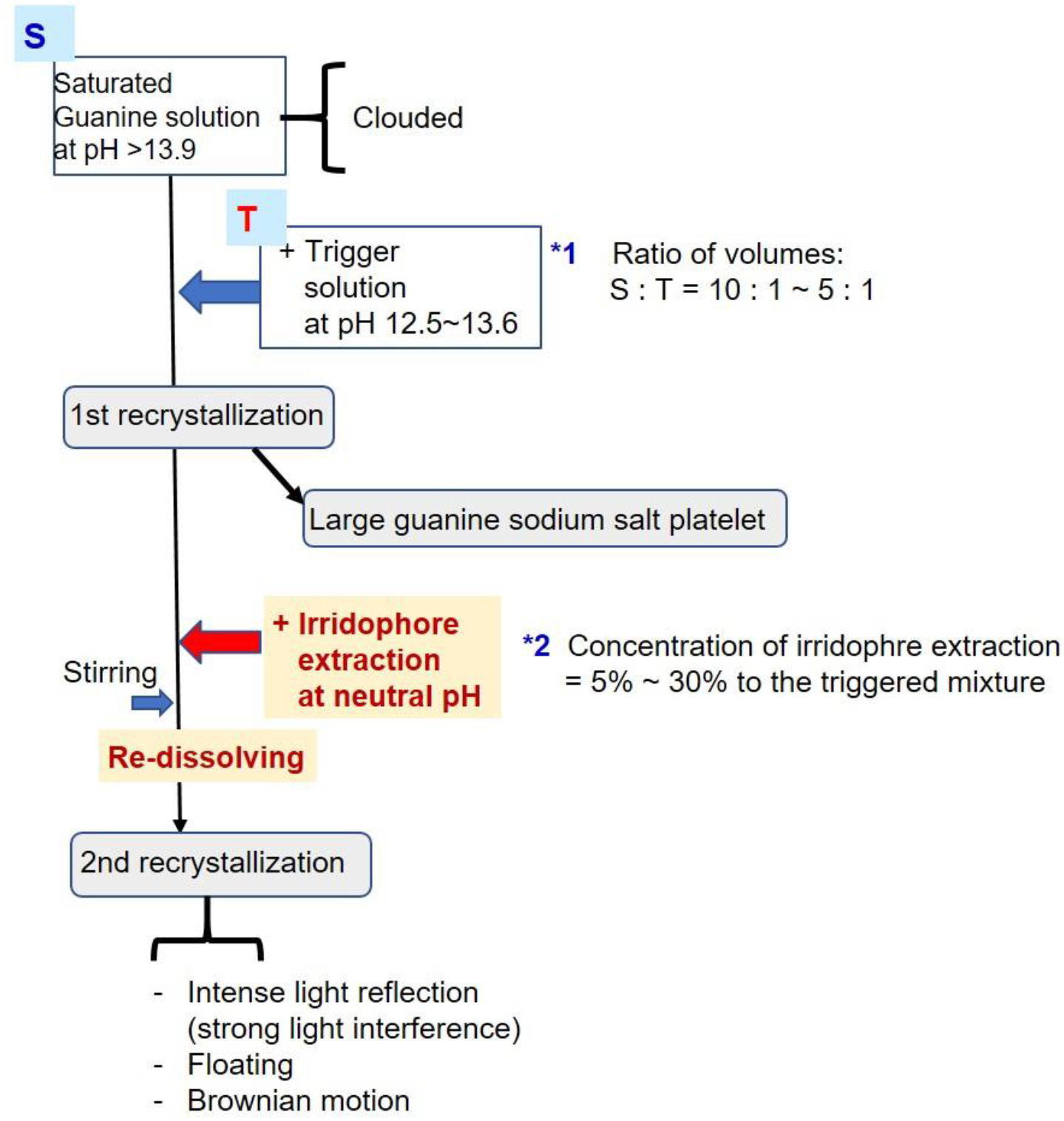
Protocols for formation of guanine sodium salt particles

**Figure 8.**
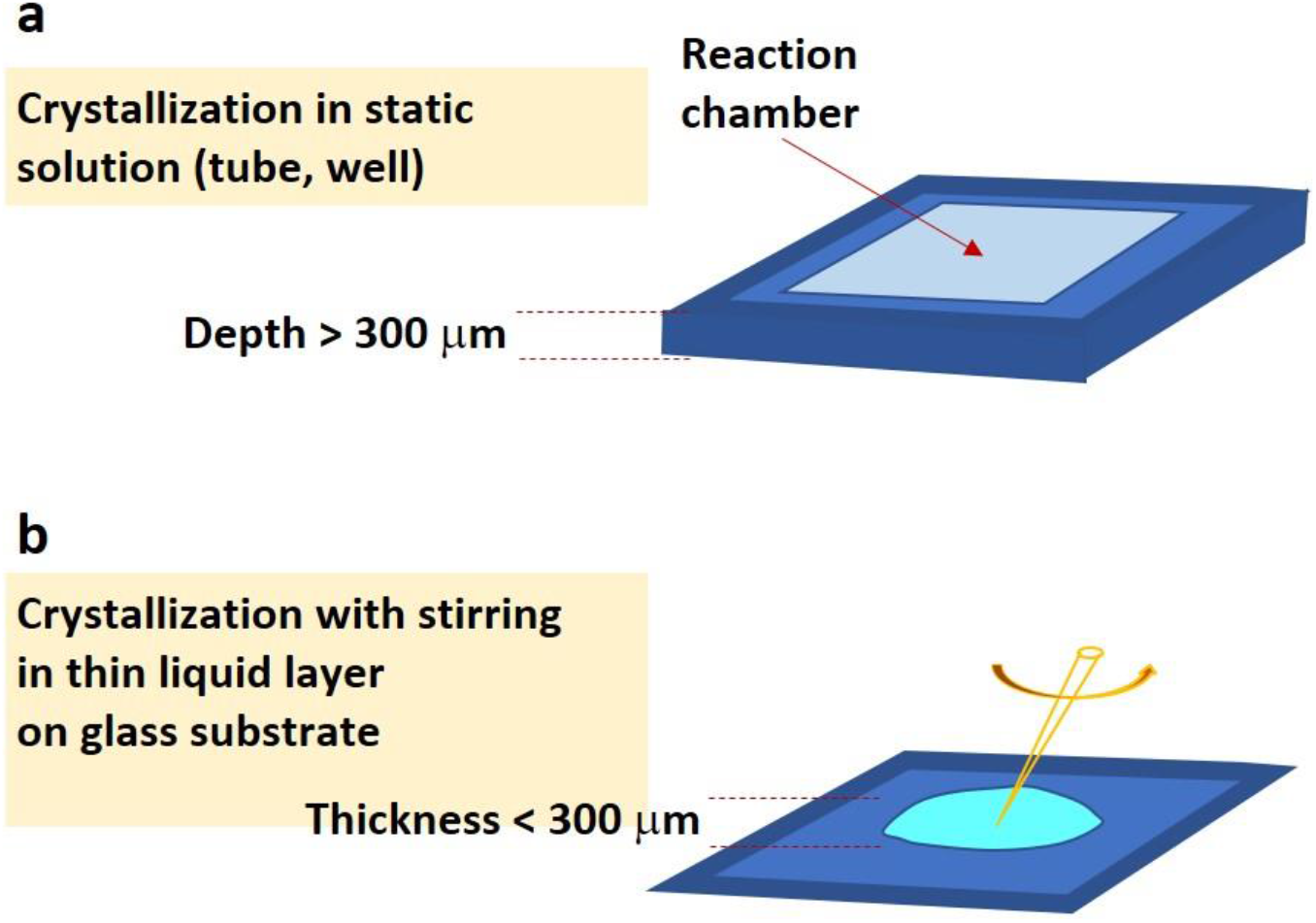
Formation of guanine sodium salt platelets in a thin layer of suspension on a glass substrate. **a**. Particles appeared slowly in the well of strong basic solution by the trigger of inserting basic solution (pH 12.5 ~ 13.6) with NaCl to condensed guanine suspension at pH>13.9. **b**. Rapid precipitation of platelet-shaped guanine sodium salt by stirring after adding the trigger solution.

So-called iridophore extraction was a mixture of major materials existed between fish scale and fish skin (explained in detail in following sub-section).

After the stimulation solution was added, the mixture was left statically (without adding mechanical stirring) in a reaction chamber, which was frame-seal incubation chamber (BIO-RAD SLF0201, USA) which is illustrated in Figure 8a or 12-mL centrifuge tube (Corning #430766, USA). The other way (Figure 8b) was employed to enhance precipitation speed. On a glass substrate (cover glass, 18mmx 18mm), the mixture forming a thin layer was stirred by a pipetting-tip (tip for max. volume of 200μl).

As mentioned in the explanation of Figure 1, after mixing IE solution with the main solution (solution S + solution T), a dissolving of both synthetic and biogenic guanine crystals occurred. Following event of precipitation provided an intensely shining particle.

### Chemical reagents

Regents of NaOH (grain, 198-13765), HCl, guanine (2-amino-6-hydroxypurine, powder, 077-01092) and distilled water were purchased from Wako Pure Chem. Ind. (Osaka Japan).

### Preparation of iridophore extraction

Iridophore-containing membrane was detached from fish scales of donor fishes. The membrane was uniformized with distilled water in a 12-mL centrifuge tube (Corning #430766) by a plastic spatula (AsOne 1-9404-02). All of the experimental methods for fish species were performed in accordance with the relevant guidelines and regulations at Hiroshima University.

### Iridophore donors

A fresh water carp species (*Cyprinus carpio*) and goldfish species (*Carassius auratus*) were utilized as an source of biogenic guanine crystals. The experimental plan using the goldfish was proposed and accepted by the bio-ethics committee at Hiroshima University (F16-2).

### Microscope, optical measurement, image analyses

A CCD microscope with 250x magnification (UK-02, Miyoshi co. Ltd., Tokyo, Japan) was utilized for microscopic observations. An LED was utilized for dark field illumination. Optical measurement of the magnetic orientation of guanine sodium salt particles was carried out using a spectrophotometer (Flame, Ocean Optics Inc. U.S.A.) to which an optical fibre 2 m in length was connected. In a cylindrical glass tube (8 mm in inner diameter), the suspension of guanine sodium salt particles was contained.

A resistive electromagnet (WS15-40-5K-MS, max field: 500 mT, Hayama, Fukushima, Japan) was used in the magnetic field exposure experiment where. the magnetic fields were switched on and off in 1-sec.

A brightness evaluation of particles in image was carried out utilizing NIH ImageJ software (for Figure 2d, Figure 5c).

## Acknowledgement

This work was supported by JST CREST JPMJCR16N1, “Development of technology integrating photonics and biology using bio-reflectors from fish.”

## Author Contribution

M.I. conceived and led the study. The paper was written by M. I.

## Additional Information

The author declares no competing interests.

## References

1. Vukusic, P & Sambles, J.R. Photonic structures in biology. Nature 424, 852–855 (2003).

2. Srinivasarao, M. Nano-optics in the biological world: Beetles, butterflies, birds, and moths. Chemical Reviews 99, 1935–1961 (1999).

3. Kinoshita, S. Yoshioka, S. Fujii, Y. & Okamoto N. Photophysics of structural color in the Morpho butterflies. Forma 17, 103–121 (2002).

4. Parker, A. R. & Martini, N. Structural colour in animals - simple to complex optics. Optics and Laser Technology 38, 315–322 (2006).

5. Anderson, T. F. & Richards, A. G. An electron microscope study of some structural colors of insects. J. Appl. Phys. 13, 748–758 (1942).

6. Kinoshita, S., Yoshioka, S & Kawagoe, K. Mechanisms of structural colour in the Morpho butterfly: cooperation of regularity and irregularity in an iridescent scale. Proc. Royal Soc. B 269, 1417–1421 (2002).

7. Zhu, D., Kinoshita, S., Cai, D. S. & Cole, J. B. Investigation of structural colors in Morpho butterflies using the nonstandard-finite-difference time-domain method: Effects of alternately stacked shelves and ridge density. Phys. Rev. E 80, 051924 (2009).

8. Berthier, S., Charron, E. & Da Silva, A. Determination of the cuticle index of the scales of the iridescent butterfly *Morpho menelaus*. Opt. Commn. 228, 349–356 (2003).

9. Plattner, L. Optical properties of the scales of Morpho rhetenor butterflies: theoretical and experimental investigation of the back-scattering of light in the visible spectrum. J. Royal Soc. Interface 1, 49–59 (2004).

10. Wickham, S., Large, M. C. J., Poladian, L. & Jermiin, L. S. Exaggeration and suppression of iridescence: the evolution of two-dimensional butterfly structural colours. J. Royal Soc. Interface 3, 99–108 (2006).

11. Fu, F., Chen, Z., Zhao, Z., Wang, H., Shang, L., Gu, Z., & Zhao Y. Bio-inspired self-healing structural color hydrogel. Proc. Natl. Acad. Sci. USA 114, 5900–5905 (2017).

12. Fu, Y., Tippets, C. A., Donev, E. U. & Lopez, R. Structural colors: from natural to artificial systems. Nanomed. Nanobiotechnol. 8, (2016).

13. Xiao, M., Li, Y., Allen, M. C., Deheyn, D. D., Yue, X., Zhao, J., Gianneschi, N. C., Shawkey, M. D. & Dhinojwala, A. Bio-Inspired structural colors produced via self-assembly of synthetic melanin nanoparticles. ACS Nano 9, 5454–5460 (2015).

14. Xu, J. & Guo, Z. Biomimetic photonic materials with tunable structural colors. J. Colloid Interface Sci. 406, 1–17 (2013).

15. Zhao, Y., Xie, Z., Gu, H., Zhua, C. & Gu, Z. Bio-inspired variable structural color materials. Chem. Soc. Rev. 41, 3297–3317 (2012).

16. Denton, E. J. Review Lecture: On the organization of reflecting surfaces in some marine animals. Philos. Trans. R. Soc. London Ser. B 258, 285–313 (1970).

17. Denton, E. J. & Land, M. F. Mechanism of reflexion in silvery layers of fish and cephalopods. Proc. R. Soc. Lond. B 178, 43–61 (1971).

18. Denton, E. J., Herring, P. J., Widder, E. A., Latz, M. F. & Case, J. F. The roles of filters in the photophores of oceanic animals and their relation to vision in the oceanic environment. Proc. R. Soc. Lond. B 225, 63–97 (1985).

19. Herring, P. J. Reflective systems in aquatic animals. Comp. Biochem. Physiol. 109A, 513–546 (1994).

20. Herring, P. J. & Cope, C. Red bioluminescence in fishes: on the suborbital photophores of *Malacosteus*, *Pachystomias* and *Aristostomias*. Mar. Biol. 148, 383–394 (2005).

21. Partridge, J. C., Douglas, R. H., Marshall, N. J., Chung, W.-S., Jordan, T. M. & Wagner, H.-J. Reflecting optics in the diverticular eye of a deep-sea barreleye fish (*Rhynchohyalus natalensis*), Proc. R. Soc. B 281, 3223 (2014).

22. Jordan, T. M., Partridge, J. C. & Roberts, N. W. Non-polarizing broadband multilayer reflectors in fish. Nature Photon. 260, 759–763 (2012).

23. Iwasaka, M & Mizukawa, Y. Light reflection control in biogenic micro-mirror by diamagnetic orientation. Langmuir 29, 4328–4334 (2013).

24. Iwasaka, M., Mizukawa, Y. & Roberts, N. W. Magnetic control of the light reflection anisotropy in a biogenic guanine microcrystal platelet, Langmuir 32, 180–187 (2016).

25. Iwasaka, M., Miyashita, Y., Mizukawa, Y., Suzuki, K., Toyota, T. & Sugawara, T. Biaxial alignment control of guanine crystals by diamagnetic orientation, Appl. Phys. Exp. 6, 037002 (2013).

26. Iwasaka, M., Miyashita, Y., Kudo, M., Kurita, S. & Owada, N. Effect of 10-T magnetic fields on structural colors in guanine crystals of fish scales, J. Appl. Phys. 111, 07B316 (2012).

27. Levy-Lior, A., Pokroy, B., Levavi-Sivan, B., Leiserowitz, L., Weiner, S. & Addadi, L. Biogenic guanine crystals from the skin of fish may be designed to enhance light reflectance. Cryst. Growth Des. 8, 507–511 (2008).

28. Gur, D., Pierantoni, M., Dov, N. E., Hirsh, A., Feldman, Y., Weiner, S. & Addadi, L. Guanine crystallization in aqueous solutions enables control over crystal size and polymorphism, Cryst. Growth Des. 16, 4975–4980 (2016).

29. Oaki, Y., Kaneko, S., & Imai, H. Morphology and orientation control of guanine crystals: a biogenic architecture and its structure mimetics, J. Mater. Chem. 22, 22686–22691 (2012).

30. Mizukawa, Y., Suzuki, K., Yamamura, S., Sugawara, Y., Sugawara, T. & Iwasaka, M. Magnetic manipulation of nucleic acid base microcrystals for DNA sensing. IEEE Trans. Magn., 50, 5001904 (1-4) (2014).

